# Temporal single-cell transcriptomic analysis of the *sox1a:eGFP* transgenic line identified the lateral floor plate progenitor cells as the origin of intraspinal serotonergic neurons

**DOI:** 10.1101/2023.03.01.530240

**Authors:** Fushun Chen, Melina Köhler, Gokhan Cucun, Masanari Takamiya, Caghan Kizil, Mehmet Ilyas Cosacak, Sepand Rastegar

**Author notes:** Author to whom correspondence should be addressed (S.R.). These authors equally contributed to this work (Mehmet Ilyas Cosacak, Sepand Rastegar).

## Abstract

The Sox family of transcription factors plays a crucial role in the development of the vertebrate nervous system. In the zebrafish embryo, *sox1* genes are expressed in neural progenitor cells and neurons of the ventral spinal cord. We recently reported that the loss of function of *sox1a* and *sox1b* leads to a significant decline in a subtype of V2 neurons, called V2s, in zebrafish. Here, a single-cell RNA sequencing approach was used to analyse the transcriptome of *sox1a* lineage progenitors and neurons in the zebrafish spinal cord at four different time points during the first five days of embryonic development, using the Tg(*sox1a:eGFP*) line. In addition to the previously described *sox1a*-expressing neurons, we found that *sox1a* is also expressed in late-developing intraspinal serotonergic neurons (ISNs). Analysis of developmental trajectories from single-cell data and depletion of lateral floor plate (LFP) cells by *nkx2*.*9* morpholino knockdown suggest that ISNs arise from LFP precursor cells. Pharmacological inhibition of the Notch signalling pathway indicates that this pathway is required for the negative regulation of the development of LFP progenitor cells into ISN populations. Our results show that the zebrafish LFP is a precursor domain that longitudinally gives rise to ISNs in addition to the previously described KA” and V3 interneurons.

## Introduction

Differential expression and combinatorial action of distinct transcription factors (TFs) are essential during embryonic development as they establish unique transcriptional expression programmes leading to tissue-specific cell fate determination (Ferg et al., 2014; Reiter et al., 2017). In the vertebrate spinal cord, dorsal and ventral antagonistic signals trigger the expression of combinations of TFs in neural progenitor domains at distinct dorsoventral (DV) positions along the spinal cord axis (Dessaud et al., 2008; Le Dreau and Marti, 2012; Sagner and Briscoe, 2019). As a result, different types of post-mitotic neurons are produced at different DV positions (Alaynick et al., 2011; Lai et al., 2016; Lu et al., 2015). In the ventral spinal cord motor neurons (MN) and ventral interneurons (IN) are specified in response to different concentrations of the morphogen Sonic hedgehog (Shh) released from the notochord and the medial floor plate (Goulding, 2009; McMahon, 2000). Close to the source of Shh, Kolmer-Agduhr (KA”) and V3 interneurons are generated. Further on, the MNs, KA’, V2, V1, and V0 interneurons are formed (Andrews et al., 2019; Dessaud et al., 2008; Yang et al., 2010; Yang et al., 2020).

There is increasing evidence that these initial subdivisions subsequently differentiate into distinct neuronal subtypes (Delile et al., 2019). For example, in the mouse embryo, the p2 progenitor domain located one level below p1 gives rise to V2 neurons. These neurons generate at least three different subtypes of V2 interneurons, called V2a, V2b, and V2c (Karunaratne et al., 2002; Li et al., 2005; Panayi et al., 2010; Smith et al., 2002; Zhou et al., 2000).

In a whole mount *in situ* hybridization gene expression screen for TFs expressed in the zebrafish spinal cord, we identified two orthologs of the mammalian *sox1* gene (Armant et al., 2013). We showed that zebrafish *sox1* genes are expressed in most ventral progenitor domains, the KA’’ and KA’ interneurons, and in a subpopulation of V2 interneurons called V2s (Gerber et al., 2019). V2s resemble mouse V2c neurons in that they express *sox1* genes and require Sox1 activity for differentiation. As in mouse V2b and V2c, in zebrafish V2b and V2s are derived from common progenitors, and V2c and V2s evolve after the formation of V2a/b subpopulations. In the absence of Sox1a and Sox1b activity, V2b/s precursors follow the V2b fate by default (Gerber et al., 2019).

We used a single-cell RNA sequencing approach to identify and analyse the transcriptome of *sox1a* lineage progenitors and neurons in the zebrafish spinal cord at 1-, 2-, 3-, and 5-days post-fertilization (dpf). Our data show that in addition to the previously described expression of *sox1a* in KA and V2s interneurons, this gene is also detected in the late developing intraspinal serotonergic neurons (ISNs). Although the role of ISNs is well studied and these neurons have been shown to contribute to the modulation of locomotor activity, axon regrowth and restoration of function after spinal cord injury in zebrafish (Barreiro-Iglesias et al., 2015; Gabriel et al., 2009; Huang et al., 2021; Montgomery et al., 2018), the origin of these neurons has not been studied. Here, the Reconstruction of the lateral floor plate (LFP) cell trajectory, the precursor of ISN (ISN-pre) and ISNs suggests that ISNs originate from the most ventral precursor domain of the spinal cord, namely the LFP. Morpholino-induced loss-of-function of *nkx2*.*9*, a TF gene crucial for the development of LFP precursor cells (Yang et al., 2010), and pharmaceutical inhibition of Notch signalling further confirmed that ISNs are derived from the LFP progenitors and that their generation requires the down-regulation of Notch signalling.

## Results

### Besides KA and V2s interneurons, *sox1a* is also expressed in ISNs and multiple neural progenitor populations in the zebrafish spinal cord

To obtain an exhaustive overview of all cell populations in the zebrafish spinal cord expressing the High Mobility Group Box *sox1a* gene and to analyse the temporal changes in the transcriptional profile of different *sox1a*-positive neural progenitors and neuronal populations, the body (trunk and tail) of the transgenic *sox1a:eGFP* zebrafish line at 1, 2, 3 and 5 dpf was manually dissected, cells dissociated and sorted GFP-positive (GFP^+^) cells subjected to single-cell RNA-sequencing (Fig. 1A). Unsupervised cell clustering by Seurat (Hao et al., 2021) on the integrated data from all four developmental stages revealed 25 unique cell clusters (Fig. 1B; Fig. S1A and B) and the characteristic marker genes expressed in the major cell types are displayed by heatmap (Fig. 1C). As the *Tg(sox1a:eGFP)* line was used to sort the spinal cord cells, we verified the co-expression of *gfp* and *sox1a* genes in each group and found that *gfp* was highly expressed in *sox1a*-positive cells (Fig. 1D), especially in cells corresponding to the lateral line and neural progenitor/interneuron (Fig. 1D). The co-expression of *gfp* and *sox1a* confirmed the enrichment of *sox1a* cells in the zebrafish spinal cord. Other cell groups identified were mesoderm, fin fold, hematopoietic system, cardiomyocytes, fast skeletal muscle, and vascular cells (Fig. 1B and C; Table S1). As previous studies have only reported *sox1a* expression in the lateral line and the neural progenitors/interneurons of the spinal cord (Gerber et al., 2019; Okuda et al., 2010), the presence of other cell groups in our analysis could be due either to the relative stability of the GFP protein and persistence of the GFP protein as possible progenitors differentiate into different cell types or the autofluorescence of certain cell populations during the cell sorting process, leading to the identification of artefactual cell clusters. Therefore, we eliminated the possible artefactual cell populations and the lateral line cell group, which are not relevant to this study. The remaining single-cell sequencing data were reanalysed using only the dataset corresponding to the neural progenitor/interneuron cell populations (Fig. 2A; Fig. S2A; Table S2). Using these remaining cell populations, we identified cell clusters corresponding to different progenitor cells, including LFP, V2b/s progenitors (V2b/s-pre) and ependymal radial glial cells (ERG) (Fig. 2A-E). We detected clusters of *sox1a*^+^ cells corresponding to KA”, KA’ and V2s neurons (Fig. 2A and C), as expected from previous studies (Andrzejczuk et al., 2018; Gerber et al., 2019), validating our approach. Interestingly, one new *sox1a* cluster representing the ISN (Montgomery et al., 2018; Montgomery et al., 2016) was also discovered (Fig. 2A, C and E; Table S2).

**Figure-1:**
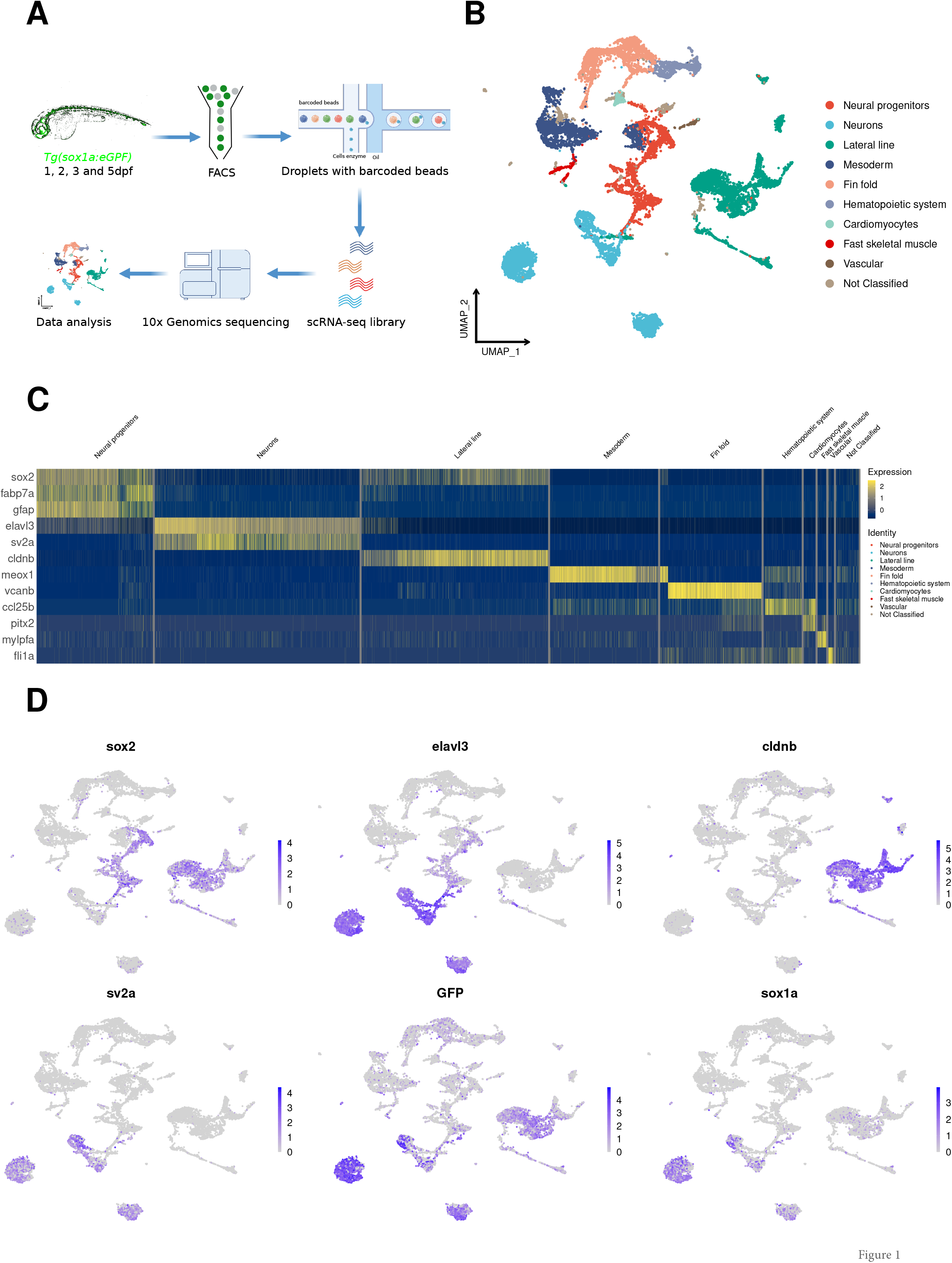
Integration of all single-cell datasets and annotation of main cell types. (A) schematic workflow of single-cell RNA sequencing. (B) UMAP shows the main cell types annotated based on the top marker and known marker genes. (C) A heatmap showing the representative marker genes expressed in main cell types. (D) Feature Plots showing marker genes used to sort *in-silico* neural progenitor and neuronal cell populations. Note: *cldnb* is used to exclude lateral line cells, which are *gfp*^*+*^.

**Figure-2:**
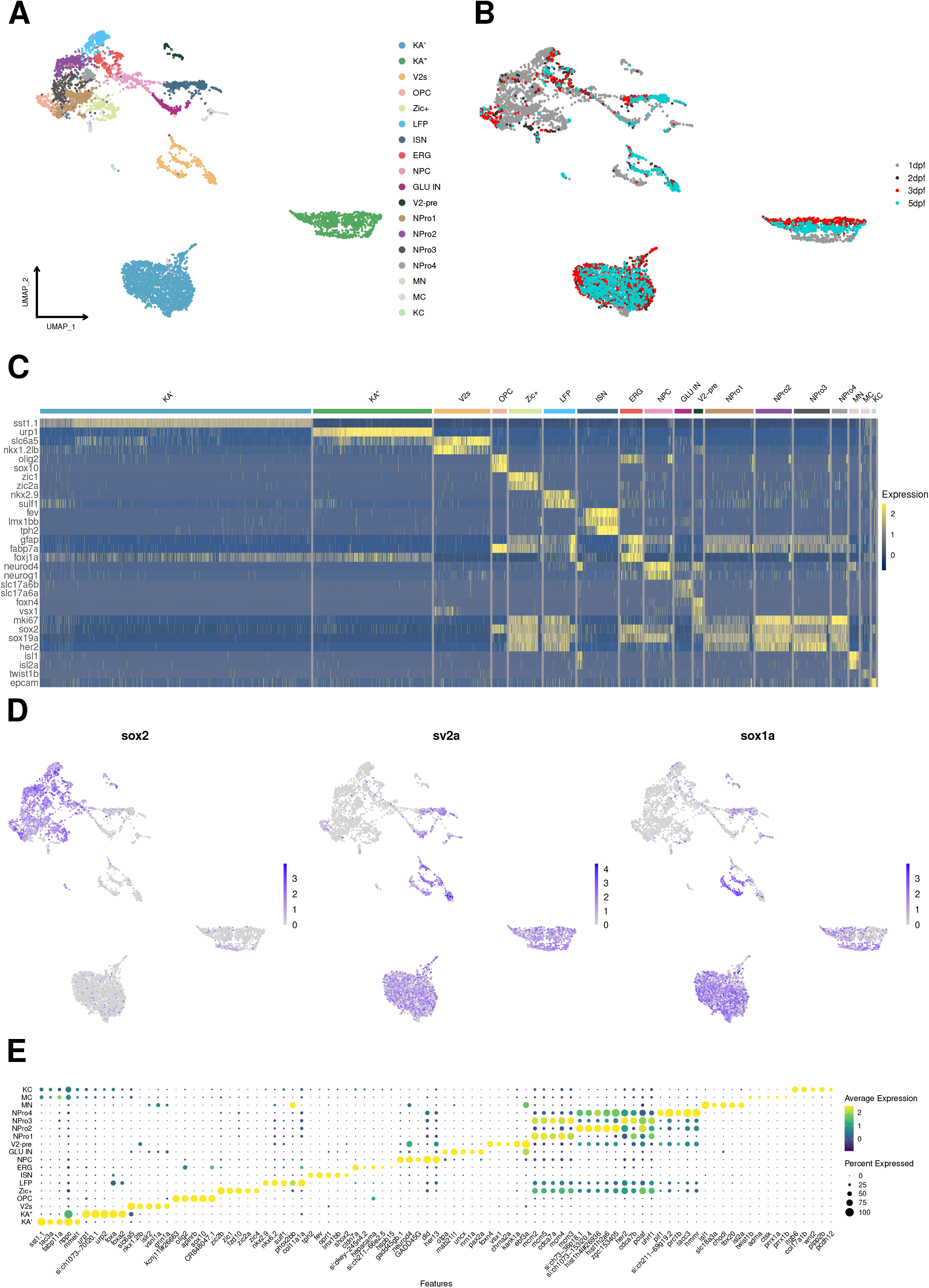
Reanalyses of the neural progenitor population and neurons identify several cell subtypes in the spinal cord. (A) UMAP shows the neural progenitors and neuronal sub-clusters. (B) The distribution of cells is based on the developmental stages. (C) A heatmap showing the representative marker genes specifically expressed in each neural progenitor and neuronal subtype. (D) Expression of *sox2* (progenitor marker) and *sv2a* (neuronal marker) on Feature Plots. (E) Dot Plot showing the top 5 marker genes for each cluster.

### The ISN cluster is made of precursor and mature neurons

To characterize the newly identified ISN cell type, we determined the genes expressed in the corresponding cell cluster (Fig. 3A; Table S2). This cell cluster is characterised by the expression of the TF genes *fev* (also known as *pet1*), *lmx1bb, gata3, gata2a, sox1a, and sox1b* as well as genes involved in the synthesis, transport, and reception of the neurotransmitter serotonin, namely *tph2* (*tryptophan hydroxylase 2*), *slc6a4a* (*solute carrier family 6 member 4*) and *htr1d* (*5-hydroxytryptamine (serotonin) receptor 1D, G protein-coupled*) (Fig. 3A; Table S2). Furthermore, hybridization chain reaction RNA fluorescent *in situ* hybridization (HCR RNA-FISH) on *Tg(sox1a:eGFP)* embryos confirmed the co-expression of GFP with *tph2* in the ventrally positioned serotonergic neurons of the zebrafish spinal cord (Fig. 3B). Temporal analysis of sequencing data indicated that these cell populations begin to emerge between two to three days of development (Fig. S3A and B). Further analysis of the data corresponding to the ISN cluster led us to subdivide this group into 3 subgroups with different and overlapping gene expression profiles (Fig. 3C-E). A cluster (cluster 1) composed mainly of cells from time points 2 and 3 dpf showed expression of genes characteristic of ISN precursor cells (ISN-pre) (Fig. 3C-E; Table S3) (Deneris and Gaspar, 2018; Deneris and Wyler, 2012). Cells in this sub-cluster were positive for *fev, gata3, gata2a, nkx2*.*2a* and *nkx2*.*2b*, but negative for *tph2* (Fig. 3E). The other two sub-clusters were both positive for *fev* and *tph2* (Fig. 3E). The largest cluster (cluster 0) is composed of cells exclusively from 5 dpf, while the smallest cluster (2) is composed of cells from 2, 3 and 5 dpf (Fig. 3D). This result demonstrates that the ISN cluster is a heterogeneous group of cells composed of a committed progenitor cluster (*fev*^*+*^ and *tph2*^*-*^ ISN-pre) and mature cluster (*fev*^*+*^ and *tph2*^*+*^ ISN).

**Figure-3:**
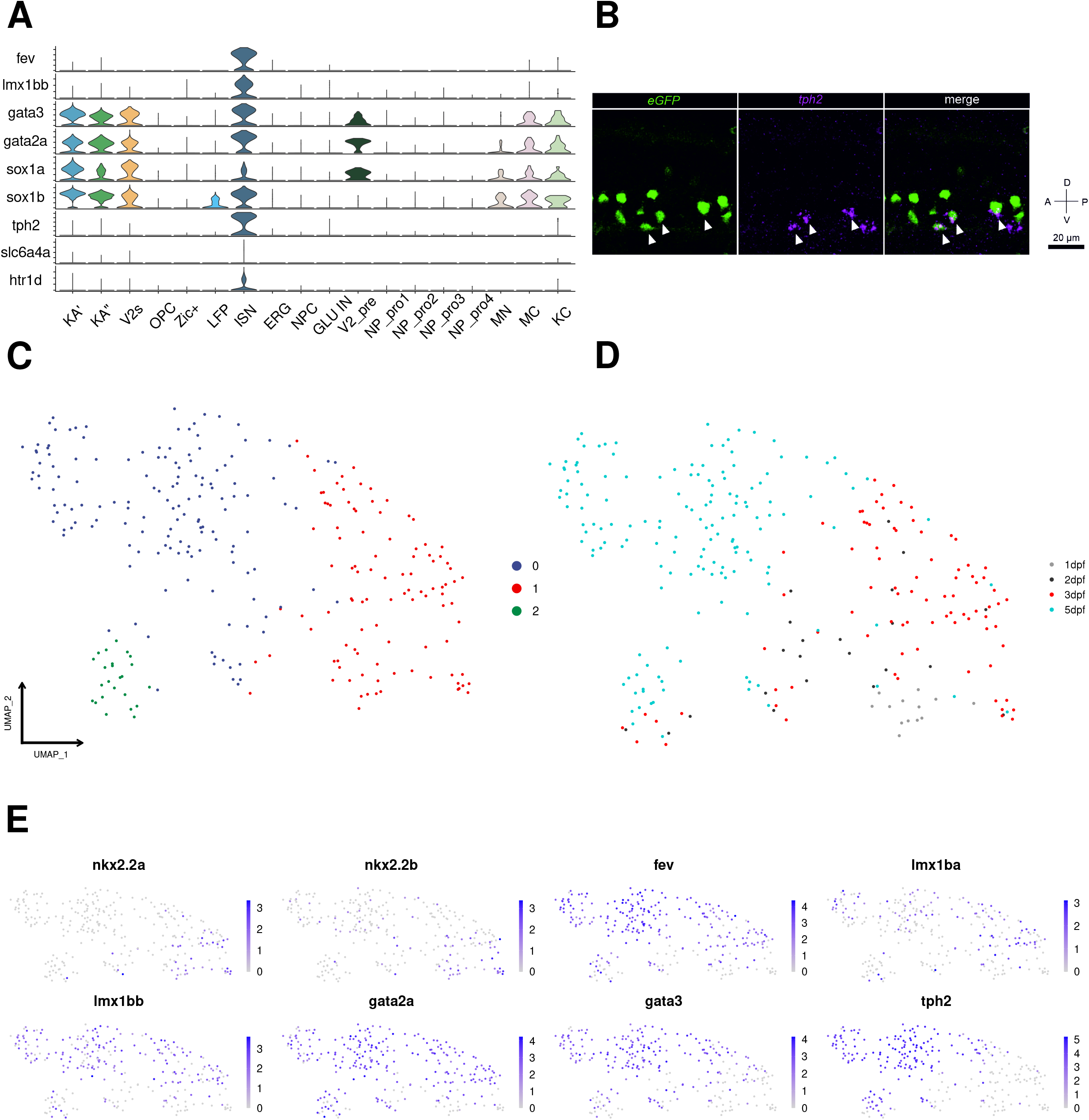
ISN cluster constitute a heterogeneous population. (A) Violin Plots showing key marker genes of ISNs co-expressed with *sox1a/b*. (B) Co-expression of *tph2 with eGFP using* HCR RNA-FISH in the zebrafish spinal cord at 3 dpf. (C) UMAP showing sub-clustering of ISN cluster into 3 sub-clusters and (D) the distribution of cells based on the developmental stages. (E) Feature Plots showing key transcription factors specifically or co-expressed in ISN-pre and ISNs.

### ISNs develop from the lateral floor plate cells

Previous cell lineage tracing and genetic data indicated that LFP progenitor cells give rise to KA” and V3 interneurons in a sequential manner (Huang et al., 2012; Jacobs and Huang, 2019; Jacobs et al., 2022; Yang et al., 2010). In this sequence of events, KA” cells are the first to be generated at about 16 hpf (hours post fertilization), followed by V3 cells at approximately 24 hpf. The specification of LFP progenitor cells into different interneuron types depends on the duration of Notch signalling. LFP cells with high Notch signalling do not differentiate and remain in a progenitor state. Inhibition of this pathway leads to the differentiation of progenitors into interneurons (Huang et al., 2012; Jacobs and Huang, 2019; Jacobs et al., 2022). Since ISNs are located ventrally in the spinal cord, the precursors of these neurons share the expression of several transcription factor genes with LFP cells such as *foxa, nkx2*.*2a* and *nkx2*.*2b* (Fig. S4A; Table S4) and in mice, some serotonergic neurons are generated from the ventral progenitor domain, which is positive for NKX2.2 and adjacent to the floor plate (Pattyn et al., 2003). We, therefore, hypothesized that in zebrafish spinal cord ISN precursor cells, are likely to originate from the LFP and that their specification, which takes place between 2 and 3 dpf (after KA” and V3), involves overlapping molecular mechanisms as for KA” and V3 interneurons.

To test this hypothesis, we re-examined the integrated single-cell data from LFP, ISN-pre and ISN clusters. Pseudotime analysis of gene expression in these clusters, where LFP was defined as the starting cluster, predicted a single lineage trajectory starting at LFP, transiting through ISN-pre, and ending at ISN (Fig. 4A). These data also clearly show that LFP cells originate from 1 dpf development stage (Fig. S4B). The ISN-pre cells are composed of 2 and 3 dpf cells while the ISNs are at the beginning of their trajectory composed of 2 and 3 dpf embryonic stage cells but at the end of the trajectory, all ISNs are from the 5 dpf developmental stage (Fig. S4B). Gene expression analysis over pseudotime shows that *nkx2*.*9* is exclusively expressed in the LFP cells (Fig. 4B; Fig. S4D; Table S4). *nkx2*.*2a* and *b* are expressed in the LFP and ISN-pre cells (Fig. 4B; Fig. S4D; Table S4). *foxa* is expressed in LFP and ISN-pre cells and some ISNs (Fig. 4B; Fig. S4D; Table S4). *lmx1bb* and *fev* marks ISN pre and ISNs cells while *tph2* is expressed mainly in ISNs (Fig. 4B; Fig. S4A and D). In line with these observations, analysis of the expression level of the *nkx2*.*9, nkx2*.*2a, foxa, lmx1bb, fev* and *tph2* genes along the developmental trajectory confirmed that *nkx2*.9 is highly expressed in most LFP cells. *tph2* is strongly expressed in ISN cells. *fev* and *nkx2*.*2a* have a broader and shallower pattern of cellular expression, both of which are co-expressed in some of the ISN-pre cells in addition to ISNs (*fev*) and LFP (*nkx2*.*2a*) (Fig. S4 D).

**Figure-4:**
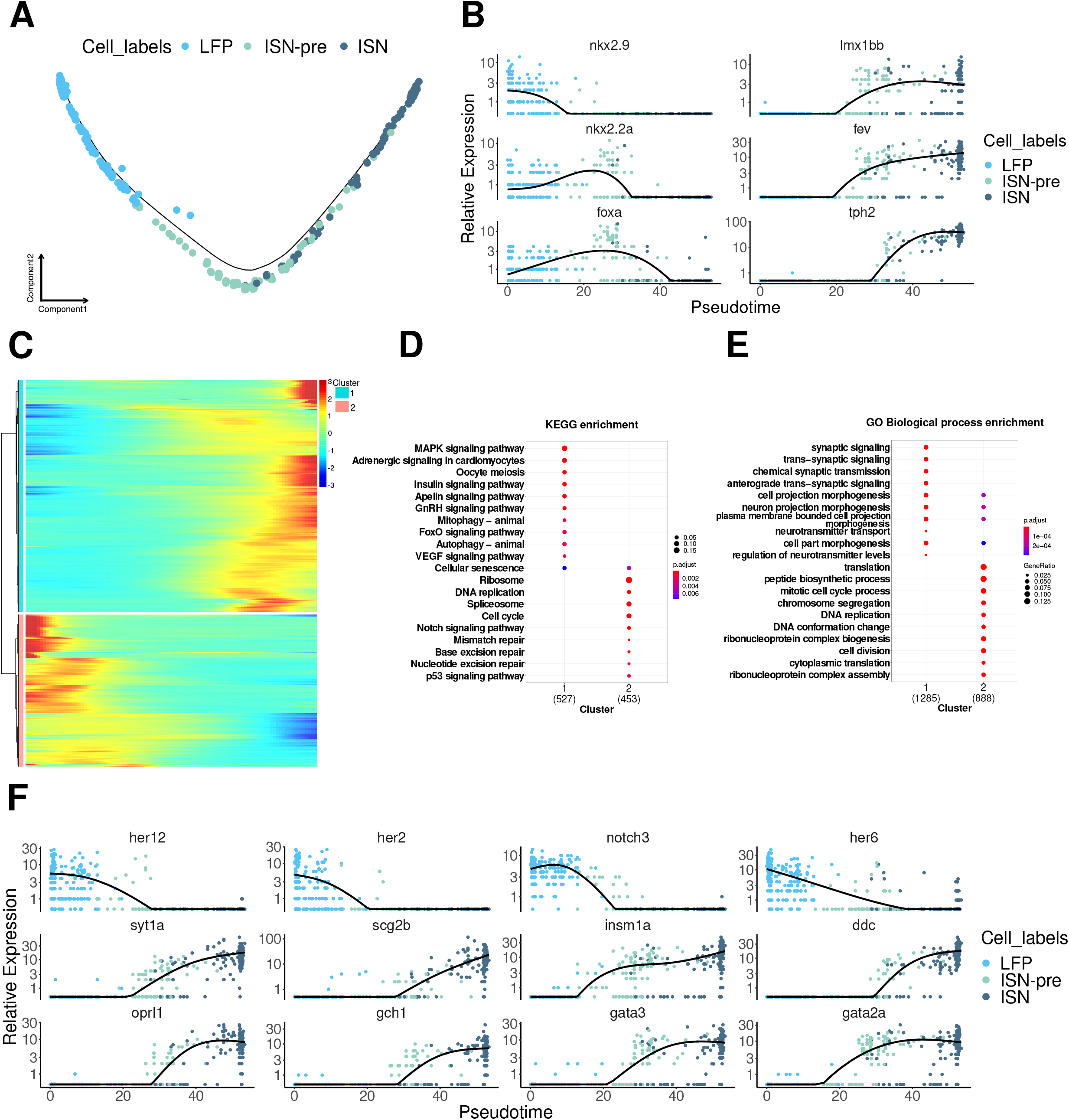
LFP is the precursor of ISN cells. (A) pseudotime cell trajectory of LFP, ISN-pre, and ISN cells. (B) The expression of key transcription markers for LFP and ISN shown on pseudotime. (C) Pseudotime heatmap showing pseudotime-dependent differentially expressed genes between clusters 1 and 2. GO and KEGG analyses of pseudotime-dependent differentially expressed genes between clusters 1 and 2., (D) GO analyses for Biological Process, (E) KEGG pathway analyses. (F) The expression of key transcription markers for LFP and ISN is shown on pseudotime.

To identify the dynamics of gene expression during the specification of LFP to ISN cells, we used monocle to identify gene modules and clusterProfiler for gene ontology (biological process) and pathway (KEGG) analyses of the module-specific genes, (Fig. 4C-4E; Table S5-7). This analysis indicated that LFP cells (cluster 2) are enriched in marker genes belonging to the Notch signalling pathway, as *her6, notch3, her12* and *her2* were among the top 25 genes of cluster 2 (Fig. 4C, D and F; Table S5-7). Other genes in the top 25 were *id1, sox3* and *mki67*, which implies that these cells are neuroprogenitors (NPro) (Table S5-7). In agreement with this result, undifferentiated LFP cells showed strong expression of Notch signalling and a decrease in this pathway triggered the differentiation of progenitors towards neuronal fate (Fig. 4F). As expected, the top 25 genes of cluster 1 (ISN, ISN-pre) were genes with a role in synaptic signalling, neurotransmitter transport or synthesis, and axonogenesis such as *synaptotagmin Ia* (*syt1a*), *dopa decarboxylase* (*ddc*), *tph2, fev, insm1a, opiate receptor-like 1* (*oprl1*), *GTP cyclohydrolase 1* (*gch1*), *gata3* and *gata2a* (Fig. 4E, C, E and F; Table S5-7).

### The integrated single-cell RNA sequencing dataset from *olig2:eGFP*^*+*^ and *sox1a:eGFP*^*+*^ spinal cord cells confirmed the common origin of V3 interneurons and ISNs

Recent work has indicated that LFP progenitors generate first KA” interneurons and later V3 interneurons (Jacobs et al., 2022). Our results suggest that ISNs, which are generated even later than V3 neurons, also develop from LFPs.

To further strengthen our findings and eventually show using single-cell RNA sequencing that both V3 and ISNs are generated from LFP cells we decided in addition to our data to analyse the single-cell RNA sequencing data published by Scott et al., (2021; Fig. S5A and B) (Scott et al., 2021). These data are of particular interest to us because these sequencing data were generated from *olig2:eGFP*^*+*^ cells of the zebrafish spinal cord. *olig2* and *sox1a* are co-expressed in many ventral cells of the spinal cord and therefore both sequencing sets share an important number of cell clusters corresponding to neuroprogenitors and neurons such as LFP, V2-pre, OPC, radial glia cells, KA” and KA’ interneurons. Noteworthy, Scott et al (2021) (Scott et al., 2021) identified two cell clusters corresponding to V3-pre (precursors) and V3 cell populations (Fig. S5 A) that were not present in our sequencing data.

We first examined the sequencing data of *olig2*^**+**^ cells at 48 hpf to identify cells potentially corresponding to ISNs. Interestingly, we were able to identify a small group of cells among the cells annotated as V3 precursors (V3-pre) that were positive for *fev, lmx1bb* and *gata2a* but negative for *tph2*, suggesting that these cells are ISN-pre cells (Fig. S 5A and B).

We then carefully analysed the cell cluster annotated as V3 in the *Olig2*^**+**^ sequencing data and found that not all cells in this cluster were *sim1a*^**+**^ (Fig. S5C), and therefore removed all *sim1a*^*-*^ cell sub-clusters from this cluster. These latter sub-clusters corresponded mainly to the *lhx5*^*+*^ group of cells (Fig. S5C). Finally, all *sim1a*^**+**^ cells, including V3-pre and V3 cells, were pooled into the LFP cluster cells and this integrated data were reanalysed (Fig. 5A and B; Fig. S5D). We identified 6 clusters corresponding to LFP, MN, NPC (neural precursor cell), ISN-pre, V3-pre and V3 cells (Fig. 5 A). The expression of genes specific to these clusters is shown in (Fig. 5B; Fig. S 5D; Table S8). In the next step, we integrated these cell clusters into the previously identified LFP, ISN-pre and ISN from our own *sox1a*^*+*^ sequencing results. The distribution and the time points origin of the corresponding cell clusters as well as the gene lists are shown in (Fig. 5C; Fig. S5E; Table S9). We observed that the first population to be generated is the V3-pre, followed by V3, ISN-pre and ISN (Fig. 5C, D; Fig. S5E).

**Figure-5:**
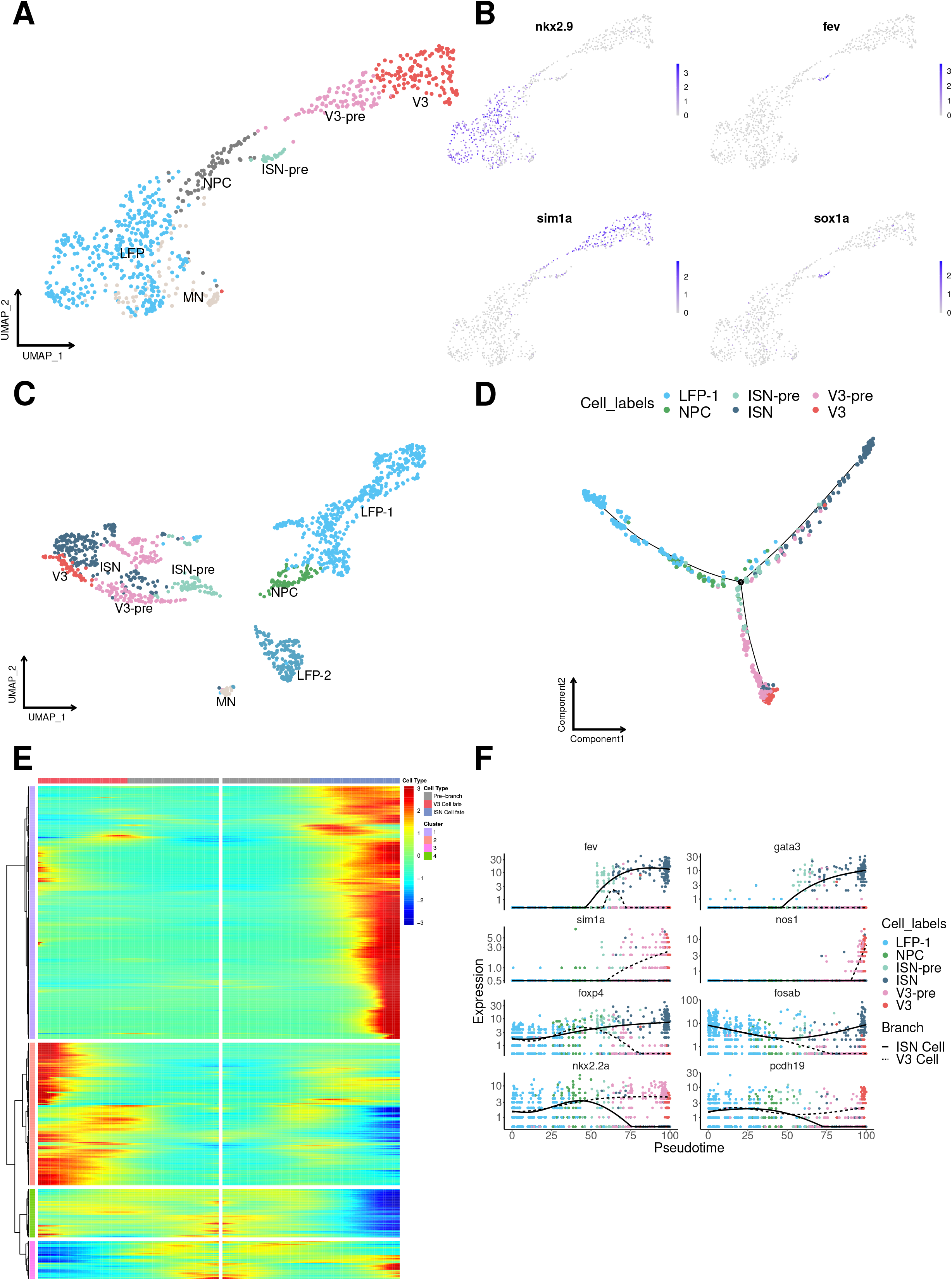
LFP is the precursor of both V3 and ISNs. (A) UMAP showing the integration of LFP, V3, and ISN cells and their subtypes (publicly available dataset; *olig2:eGFP* (Scott et al., 2021)). (B) Feature plots of marker genes for LFP and ISN cells. (C) UMAP showing the integration of LFP, V3, and ISN cells and their subtypes (current study: *sox1a:eGFP*) integrated with the same cells from *olig2:eGFP*. (D) Cell trajectory showing LFP as the precursor of V3 and ISN, (E) heatmap showing pseudo-dependent differentially expressed genes between LFP and V3/ISN cells. (F) The expression of key transcription factors for LFP and ISN/V3 is shown in pseudotime.

The developmental trajectory of the cells over pseudotime beautifully recognized the LFP cells as the origin of both neuronal populations (Fig. 5D). This trajectory from the LFP transit over NPCs further split into two branches. The lower branches of the younger cell population correspond to V3-pre and V3, and the upper branch of the older cell population contains the ISN-pre and ISN cells (Fig. 5D; Fig. S5F and G). To analyse the marker genes playing a role in this transition, we identified 4 gene clusters by monocle based on pseudotime (Fig. 5E).

From the heatmap, cluster 1 genes are highly expressed in ISNs. Their cell type-specific markers are unique to differentiated ISNs (e.g. *fev, gata3*, Fig. 5F; Table S10). Cluster 2 genes are significantly expressed in V3 cells (e.g. *sim1a, nos1*, Fig. 5F; Table S10). Cluster 3 genes are strongly expressed in LFP and ISNs during the transition to V3 and ISNs (e.g. *foxp4, fosab*, Fig. 5F; Table S10). Cluster 4 genes are specifically expressed in LFP and V3 cells during the transition to V3 and ISNs (e.g. *nkx2*.*2a, pcdh19*, Fig. 5F; Table S10).

### Depletion of LFP progenitor cells by *nkx2*.*9* morpholino knockdown results in a reduction in the number of *tph2*^+^ neurons

We have shown that V3 and ISN cells have a common origin and that both cell populations are derived from the LFP. To further support this claim, we hypothesized that functional knockdown of upstream LFP progenitor transcriptional programs would result in a reduction of serotonergic cells. Our gene expression analysis showed that among the TF markers of the LFP, the *nkx2*.*9* gene is the best candidate because this gene is exclusively expressed in the LFP while *nkx2*.*2a and nkx2*.*2b* are also expressed in ISN-pre and V3-pre as well as in V3 (Fig. 5A and 6A). Furthermore, the *nkx2*.*9* gene is a direct target of Shh that is released from the notochord and floor plate and both genes play a central role in the development of the lateral floor plate (Odenthal et al., 2000; Xu et al., 2006; Yang et al., 2010). In zebrafish, knockdown of the *nkx2* genes resulted in a sharp reduction in the number of KA” and V3 interneurons, with a consequent decrease in the expression of the markers *tal2* and *sim1a*, respectively, indicating impaired LFP development (Yang et al., 2010). We tested whether *nkx2*.*9* is required for *tph2* expression in ISNs by injecting an antisense morpholino (Mo-nkx2.9) directed to the translation start site (see Materials and Methods). The knockdown of *nkx2*.*9* translation resulted in a ∼73% reduction in *urp1*-positive cells in the lateral floor plate at 24 hpf (Fig. 6D and E) when compared to uninjected embryos (Fig. 6B and E) or control morpholino (Fig. 6C and E).

**Figure-6:**
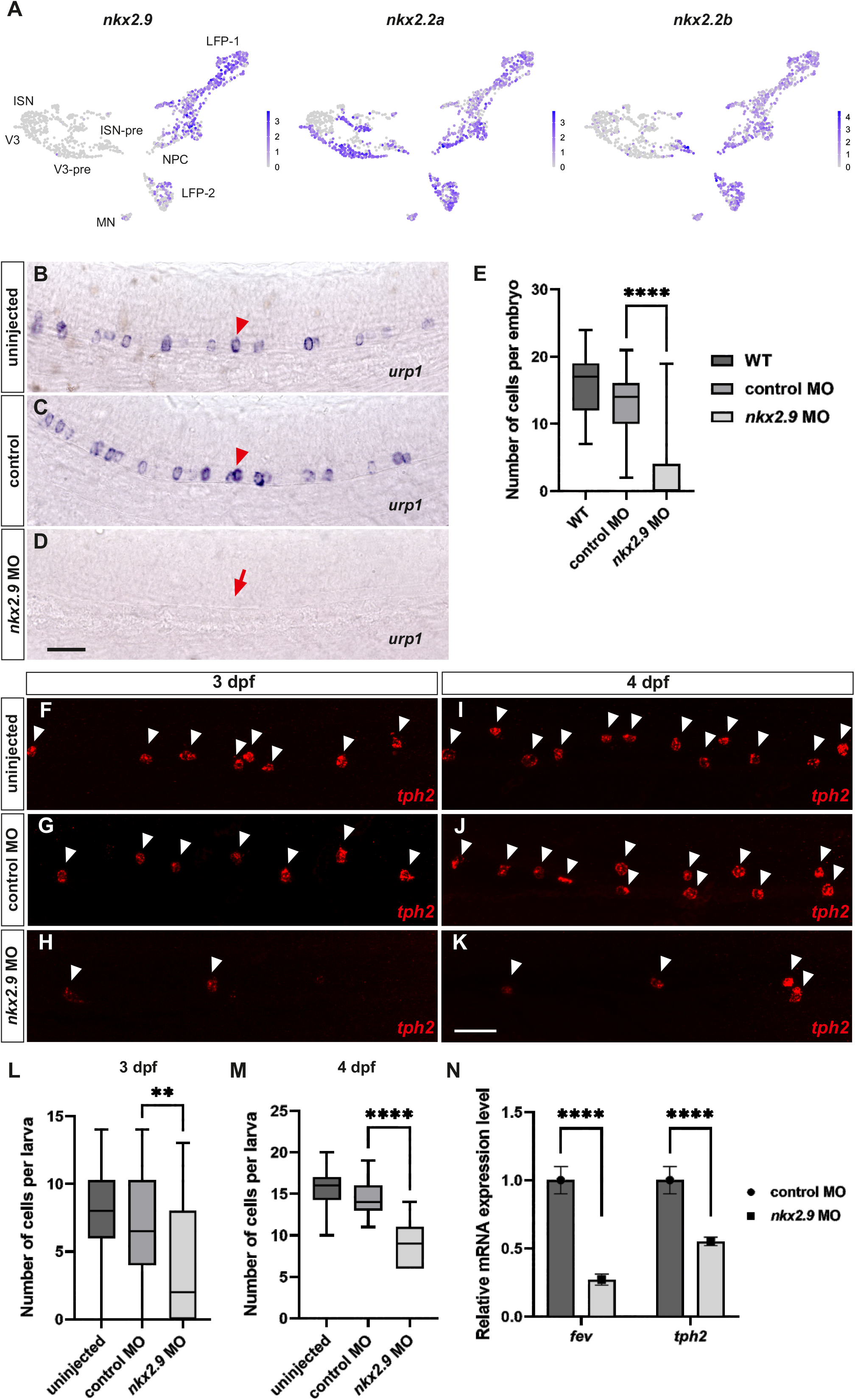
Intraspinal serotonergic neurons originate from the lateral floor plate. (A-C) Uninjected embryos (A) and embryos injected with control morpholino (B) or morpholino directed against *nkx2*.*9* mRNA (C) were hybridized to *urp1* probes at 24 hpf. Note the absence of *urp1*^+^ in *nkx2*.*9* morpholino-injected embryo in (C; red arrow). Red arrowheads indicate *urp1*^+^ cells in (A) and (B). (D) Cells were counted from the 8^th^ to the 16^th^ somite along the spinal cord from 15 to 31 embryos. Knockdown of *nkx2*.*9* resulted in a reduction of about 73 % in *urp1*-expressing cells. (E-J) Uninjected larvae (E, H) and larvae injected with control morpholino (F, I) or morpholino directed against *nkx2*.*9* mRNA (G, J) were hybridized to *tph2* HCR RNA-FISH probes at 3 and 4 dpf. (K) Cells were counted above the yolk extension over a five-somite span from 34 to 38 embryos (3 dpf) or 13 to 16 embryos (4 dpf). Knockdown of *nkx2*.*9* led to a decrease of *tph2*-positive cells by 41% and 38% at 3 and 4 dpf, respectively. (D, K, L) Statistical significance was assessed using Welch’s *t*-test, and the significance level is graded with the number of asterisks: ** for *p*<0.01, **** for *p*<0.0001. Scale bar: 25 µm. (M) RT-qPCR of *fev* and *tph2* mRNA levels of control morpholino and *nkx2*.*9* morpholino-injected larvae at 3 dpf. *Fev* and *tph2* expression is downregulated in larvae injected with *nkx2*.*9* morpholino at 3 dpf. Statistical significance was assessed using unpaired two-tailed Student’s *t*-test, and the significance level is graded with the number of asterisks: **** for *p*<0.0001. *n* = 70 embryos/condition. Abbreviations: dpf, days post fertilization.

In agreement with our hypothesis, sibling-injected embryos showed a marked reduction of 41% and 38% in *tph2*-positive cells at 3 and 4 dpf, respectively (Fig. 6F-K). The specificity of the knockdowns was controlled by the injection of a control morpholino (Fig. 6C, G and J). Quantification of the morpholino injection experiments was confirmed by real-time quantitative PCR (RT-qPCR) (Fig. 6N). The difference in percentage reduction observed between *urp1* and *tph2* in this experiment can be explained by a gradual dilution of the morpholino concentration during the development of the injected embryos. In general, morpholino injected at the 1-cell stage has a knockdown efficiency of 3 days post-injection (Bill et al., 2009).

### Notch signalling controls the differentiation of LFP cells into ISNs

Our single-cell RNA sequencing results show that LFP cells are proliferating neuroprogenitor cells as they are positive for *sox3, sox2, sox19a sox19 b* and *gfap* (Table S2). These cells also express genes involved in cell cycles such as *mki67, mcm5* and *mcm2* (Table S2). In addition, these cells strongly express genes involved in Notch signal reception and processing such as *notch3* receptor and downstream targets like *her12, her2, her9, her6, her8a* and *her4* (Table S 2).Differentiation of KA” and V3 interneurons from LFP progenitors has been shown to require downregulation of Notch signalling (Jacobs and Huang, 2019; Jacobs et al., 2022). Thus, we asked whether the development of ISNs from LFP cells also depends on the down-regulation of this pathway. We, therefore, exposed WT embryos to the γ-secretase inhibitor LY411575, which blocks Notch processing after the ligand binding (Fauq et al., 2007). We determined the number of *tph2*^*+*^ neurons in the zebrafish trunk at 3 dpf embryos exposed to LY411575 from 2 until 3 dpf (Fig. 7A). We decided to do the treatment at 2 dpf because this is when the first ISN-pre and ISN are observed in the ventral position of the spinal cord, while V3 neurons are already specified. Blocking Notch signalling in the time window between 2 and 3 dpf resulted in an important increase in the number of serotonin-positive cells compared to sibling embryos treated with DMSO alone (Fig. 7A-C). Thus, the specification of ISN cells depends on Notch signalling as shown for KA” and V3 interneurons (Jacobs et al., 2022).

**Figure-7:**
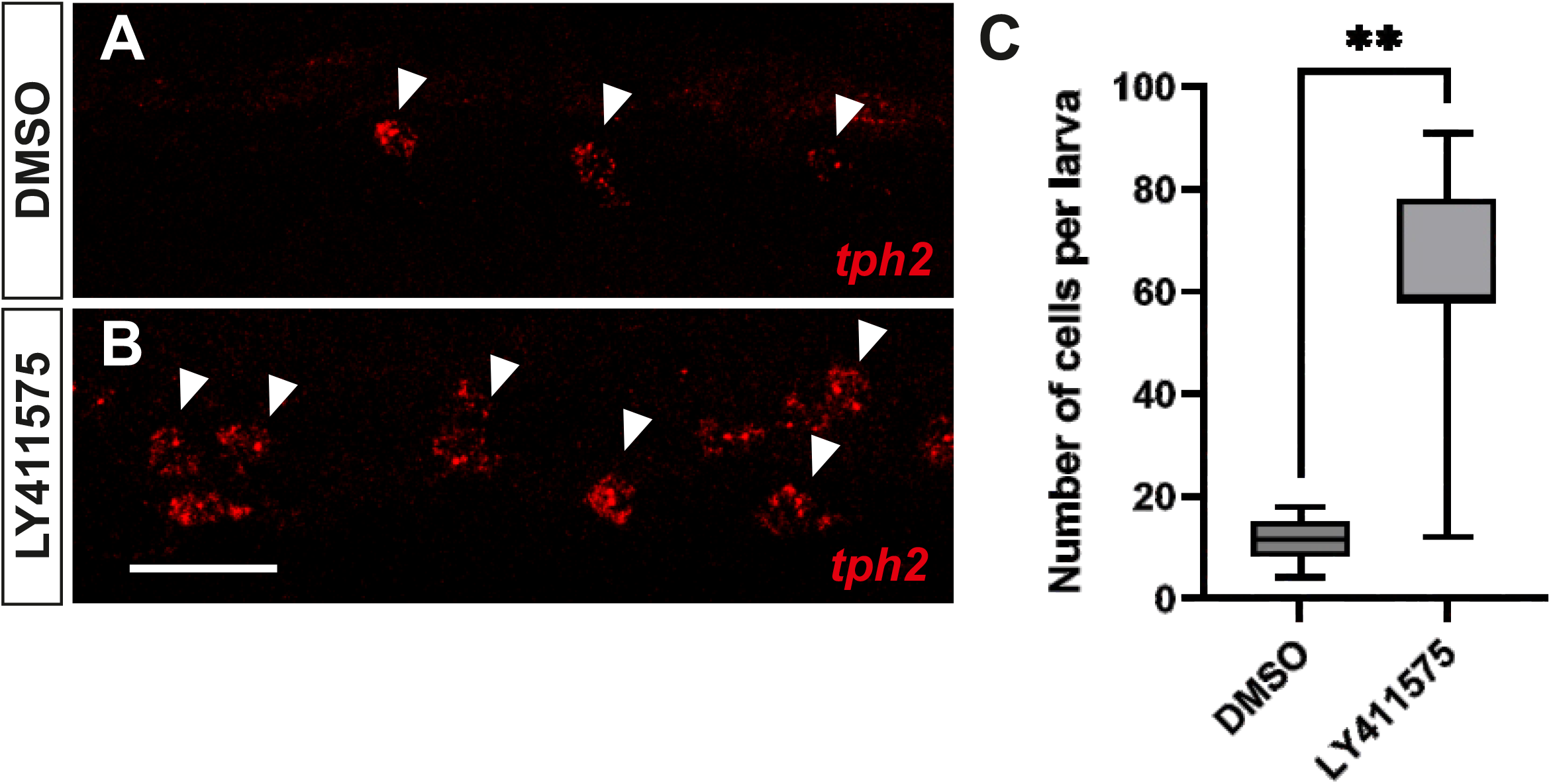
Inhibition of the notch signalling pathway leads to an increase in tph2 neurons. (A-B) HCR RNA-FISH against *tph2* mRNA. Arrowheads indicate *tph2*-expressing neurons. (A) Control embryo treated with DMSO. (B) Embryo treated with the notch inhibitor LY. (C) Quantification of *tph2*^+^ neurons. Embryos were treated from 2 to 3 dpf. Dorsal is upwards; anterior is leftwards. Statistical significance was assessed using Welch’s *t*-test, and the significance level is graded with the number of asterisks: ** for *p*<0.01. Scale bar: 20 µm.

## Discussion

Our sequencing data identified previously unidentified expression domains for *sox1a*. Here we show that in addition to KA and V2 neurons, *sox1a* is also expressed in intraspinal serotonergic neurons in the zebrafish ventral spinal cord. Our data indicate that, like KA” and V3 interneurons, ISNs develop from LFP progenitor cells through a latent specification after 48 hpf that succeeds other interneuronal subtypes. Specification of ISNs relies on down-regulation of the Notch signalling pathway like other interneurons.

### The lateral floor is the most ventral progenitor domain of the zebrafish spinal cord that encompasses the P3 domain

Until recently, the p3 progenitor domain was described as the most ventral progenitor domain of the vertebrate spinal cord from which different classes of V3 subpopulations are generated over time (Delile et al., 2019; Dessaud et al., 2008). Our single-cell RNA sequencing results and recent work based on classical lineage tracing in the zebrafish embryo (Jacobs et al., 2022) have independently demonstrated that the LFP is the most ventral progenitor domain of the spinal cord, from which three distinct ventral neural subtypes emanate at different developmental stages of the zebrafish embryo.

The p3 domain is included in the lateral floor LFP and constitutes only one of three cell progenitor populations along with the progenitor cells that give rise to the KA”, V3 and ISN interneurons. This suggests that the LFP, as a heterogeneous population of progenitor cells, could temporally give rise to neuronal subpopulations in the spinal cord and could be a principle of vertebrate-wide development. This hypothesis still needs to be studied and confirmed in other model organisms.

### The mechanisms of neuron generation from the LFP and other more dorsally positioned progenitor domains are not totally similar

In vertebrates, different populations of neurons are generated along the dorsoventral axis of the spinal cord because of the antagonistic interaction between ventral and dorsal signals. Consequently, different progenitor domains are created along this axis from which post-mitotic neurons will be generated. However, it is well accepted that the initial progenitor domains subsequently differentiate into distinct neuronal subtypes. For example, we and others have shown that in zebrafish, the p2 domain gives rise to several progenitor subdomains from which at least three V2 subtypes are derived that express distinct neurotransmitters and connect to different motoneurons (Batista et al., 2008; Bello-Rojas and Bagnall, 2022; Gerber et al., 2019; Kimura et al., 2008). Interestingly, this pattern of neuron generation and specification appears to be different in the LFP. In the latter domain, the cyclic and repetitive activation and deactivation of Notch signalling allow the generation and sequential specification of 3 different populations of interneurons from 16 hpf to 120 hpf. Data obtained in mice suggests that V3 neurons are not a homogeneous population but are composed of 4 subpopulations named V3.1, V3.2, V3.3 and V3.4 (Delile et al., 2019). These neurons probably originate from a further specification of p3 to different subtypes as has been shown for the mouse and zebrafish p2 population (Batista et al., 2008; Bello-Rojas and Bagnall, 2022; Gerber et al., 2019; Karunaratne et al., 2002; Li et al., 2005; Panayi et al., 2010; Smith et al., 2002; Zhou et al., 2000). It appears that the specification of LFP neurons combines mechanisms unique to this domain (cyclic Notch activation and deactivation) in addition to existing mechanisms (classical generation of neuronal subpopulations from progenitor cells) in other domains to generate a larger number of different neurons compared to all other spinal cord progenitor domains. The triggering of sequential activation and deactivation of Notch signalling in the different progenitor cells of the LFP still needs to be documented, as well as the existence of the same mechanisms in other animal models.

### Gene regulatory networks governing the development of serotoninergic neurons are conserved during evolution

In the mouse ventral hindbrain, where the signalling cascade involved in the specification of 5-HT neurons is most extensively studied, cells adjacent to the floor plate and positive for Nkx2.2 were shown to develop into serotonergic neurons (Pattyn et al., 2003). In addition, Nkx2.2 and Foxa2 were shown to be upstream of Gata2 and Gata3 (Jacob et al., 2007). Activation of these latter TF factors in turn induces the expression of Lmx1b and Pet1 (Craven et al., 2004) which leads to the expression of 5-HT in serotonergic neurons (Hendricks et al., 1999; Hendricks et al., 2003). Consistent with this cascade of gene expression, the binding sites of Gata and Pet1 on the promoter regions of Pet1 and Tph2, respectively, were identified and found to be essential for the expression of these genes in serotonergic neurons (Krueger and Deneris, 2008; Scott et al., 2005; Wyler et al., 2016). Genetic pieces of evidence indicate a pivotal role of Gata2 in the generation of 5-HT neurons (Craven et al., 2004). Our single-cell RNA sequencing suggests similar gene regulatory networks involved in the development of ISN neurons in zebrafish. The cascade of gene regulation during the specification of ISN in zebrafish is well conserved during evolution because it involves genes and anatomical regions almost identical to those in the mouse embryo. As pre-ISN cells appear slightly earlier than ISN neurons during zebrafish spinal cord development (Fig. S4B), these cells are postmitotic (*elavl3*^+^) and express *fev, lmx1bb, lmx1ba, nkx2*.*2a, nkx2*.*2b, gata2a* and *foxa*, and only slightly *tph2*^*+*^, unlike ISNs which are strongly expressing *tph2* (Fig. S4A).

## Acknowledgements

We thank T. Beil, A. Schröck and M. Sobucki for technical support; C. Lillesaar for sharing the zebrafish *pet1:eGFP* line (Lillesaar et al., 2007). We are deeply grateful to U. Strähle for his financial support in the initial phase of this project.

## Author contributions

Conceptualization: S.R., Methodology: C.F, M.K, G.C, M.T, M.I.C; Software: C.F, M.I.C; Validation: C.F, M.K, G.C, M.T, M.I.C; Formal analysis: C.F, M.K, G.C, M.T, M.I.C; Investigation: C.F, M.K, G.C, M.T, M.I.C; Resources: S.R, M.I.C, C.K Data curation: S.R., C.F, M.I.C; Writing - original draft: S.R., M.I.C, C.K Writing - review & editing: S.R., C.K; Visualization: C.F, M.K, G.C, M.T, M.I.C; Supervision: S.R. Project administration: S.R. Funding acquisition: S.R.

## Data availability

Single-cell RNA sequencing data of the *sox1a:eGFP* line have been deposited in NCBI’s Sequence Read Archive (SRA) under accession number <u>XXX</u>.

## Ethics statement

The permit for the operation of animal husbandry has been obtained according to §11 TSchG (Regierungspräsidium Karlsruhe, Aktenzeichen 35-9185.64/BH KIT). The animals are kept according to the recommendations of the European Society for Fish Models in Biology and Medicine (EuFishBioMed) of the Karlsruhe Institute of Technology (KIT), and according to the respective white paper (Alestrom et al., 2020). All experiments with zebrafish embryos were done in developmental stages between 24 hpf and 120 hpf, zebrafish embryos are not protected during this period (Strahle et al., 2012).

## Materials and Methods

### Fish stocks and embryo collection

Wild-type zebrafish were obtained from the ABO line, an inbred line initially derived from an intercross between the AB and OX lines (European Zebrafish Resource Centre). Wild-type and transgenic zebrafish *Tg(sox1a:eGFP)* (Gerber et al., 2019), *Tg(pet1:eGFP)* (Lillesaar et al., 2007) were maintained on a 14 h/10 h light-dark cycle at 28.5°C in a recirculation system (Schwarz) and fed commercial food and in-house hatched brine shrimp as described previously (Westerfield, 2000). Embryos were cultured in an embryo medium and staged according to Kimmel et al. (1995) (Kimmel et al., 1995).

For the single-cell sequencing, *Tg(sox1a:eGFP)* and WT-AB fish were crossed and embryos were collected within 15 min to minimize the age difference between embryos. Dead and unfertilized eggs were removed at 4 hpf. Batches of 40 embryos were collected and kept in 30 ml of E3 zebrafish medium with methylene blue at 28°C and in a dark incubator until they reached the desired developmental stage.

### Cell Dissociation

Cell dissociation was performed as described (Cosacak et al., 2020; Cosacak et al., 2019). A total of 60 embryos at developmental stages 1, 2, 3 and 5 dpf were kept on ice for 10 min, then the trunk was separated from the head and the yolk removed. The bodies were placed directly into the dissociation buffer and kept on ice. Dissociation was done at 28°C by rotating the cells and trituration was done every 10 min until complete dissociation. Cells were filtered through 40 µm filters (Falcon, Cat # 431750) and centrifuged at 300 g in 2% BSA and resuspended in 4% BSA. A total of 22,000 cells were sorted based on the GFP signal and the cells were subsequently treated for encapsulation.

### Single-cell Transcriptomics

Single-cell transcriptome sequencing was performed based on the 10x Genomics Single-cell transcriptome workflow (Zheng et al., 2017). Specifically, 22,000 FACS-sorted cells were recovered in BSA-coated tubes containing 5 µl PBS. Accurate cell numbers were obtained using a Neubauer Hemocytometer. Cell samples were then carefully mixed with Reverse Transcription Reagent (Chromium Single Cell NextGem 3’ Library Kit v3.1) and loaded onto a Chromium Single Cell G Chip to reach a recovery of up to 10,000 cells per sample. The samples were processed further following the 10x Genomics user manual guidelines for single cell 3’ RNA-seq v3.1. In short, the droplets were directly subjected to reverse transcription, the emulsion was broken, and cDNA was purified using silane beads (Chromium Single Cell 3’ Gel Bead Kit v2). After amplification of cDNA with 11 cycles, samples were purified with 0.6x volume of SPRI select beads to deplete DNA fragments smaller than 400 bp and cDNA quality was monitored using the Agilent FragmentAnalyzer 5200 (NGS Fragment Kit). 10 µl of the resulting cDNA was used to prepare single-cell RNA-seq libraries - involving fragmentation, dA-Tailing, adapter ligation and 10-13 cycles of indexing PCR following manufacturers guidelines. After quantification, libraries were sequenced on multiple Illumina NovaSeq 6000 S4 flowcells in 100bp paired-end mode, aiming for 30,000 fragments per cell. The raw sequencing data were processed with the ‘count’ command of the Cell Ranger software (v6.1.2) provided by 10X Genomics. To build the reference, the zebrafish genome (GRCz11), as well as gene annotation (Ensembl 104), were downloaded from Ensembl and the annotation was filtered with the ‘mkgtf’ command of Cell Ranger to include gene of the following types: ‘protein_coding’, ‘lincRNA’, ‘antisense’, ‘IG_LV_gene’, ‘IG_V_gene’, ‘IG_V_pseudogene’, ‘IG_D_gene’, ‘IG_J_gene’, ‘IG_J_pseudogene’, ‘IG_C_gene’, ‘IG_C_pseudogene’, ‘TR_V_gene’, ‘TR_V_pseudogene’, ‘TR_D_gene’, ‘TR_J_gene’, ‘TR_J_pseudogene’, ‘TR_C_gene’. Sequence and annotation of the GFP construct were included manually. Genome sequence and filtered annotation were then used as input to the ‘mkref’ command of Cell Ranger to build the appropriate Cell Ranger Reference.

### Seurat Clustering and Integration of datasets

We used Seurat (version 4) (Hao et al., 2021) and the following common steps for each round of analysis. Cells (i) with more than 20% mitochondrial RNA genes, (ii) with less than 500 total reads (nCount_RNA), and 200 unique genes (nFeature_RNA) were removed from analyses. Genes found in less than 3 cells were also removed from analyses. The remaining cells were converted to Seurat objects, data was normalized (NormalizeData), the top 2000 variable genes were identified (FindVariableFeatures), and all genes were used to scale data (ScaleData) by regressing out nCount_RNA. The top 30 PCAs (RunPCA) were calculated and these PCAs were used for dimensional reduction (RunUMAP), finding cell clusters (FindNeighbors, FindClusters) with resolution = 0.5. To integrate all datasets, we used Seurat objects and the top 2000 variable genes identified above. First, the anchors were identified (FindIntegrationAnchors), objects were integrated (IntegrateData), and all genes were used for scaling data (ScaleData) by regressing out nCount_RNA. The top 30 PCAs (RunPCA) was calculated and these PCAs were used for dimensional reduction (RunUMAP), finding cell clusters (FindNeighbors, FindClusters) with resolutions 0.5, 1, 1.5 and 2. Unless indicated the above steps and settings were used for sub-setting cells for iterative analyses.

### Preliminary Quality Control

A preliminary analysis of 1 dpf embryos showed a cell population that is highly expressing *hbb* genes (e.g., *hbbe1*.*1*). We used scaled data from 1 dpf embryos Seurat object and labelled cells that are expressing the *hbbe1*.*1* gene more than 0.0. These cells were removed from 1 dpf embryos and another round of clustering was performed as above for 1 dpf raw counts (*hbb* cells removed). The remaining *hbb* cells, which were clustered together, were removed as well. After quality control and removing *hbb* expressing cells from the 1 dpf dataset, developmental stages 1, 2, 3 and 5 dpf datasets were converted to a Seurat object, and the above settings were used for all datasets.

### Identifying Main Cell Types

After integration and clustering, we identified 24 and 40 clusters of cells using the resolutions 0.5 and 1.5, respectively. We identified the marker genes using the FindAllMarkers function. We annotated the main cell types using the top 20 marker genes and known marker genes from the literature. For example, we used *fabp7*a/*sox2*/*gfap* for neuralprogenitors (NPro), *elavl3*/*sv2a* for neurons, *cldnb*/*epcam*/*sox2* for lateral lines, *meox1* for mesoderm, *vcanb* for fin fold, *ccl25b* for the hematopoietic system, *pitx2* for cardiomyocytes, *mylpfa* for skeletal muscle and fli1a for the vascular system. The remaining cells without clear markers were labelled as “unclassified” cells.

### Iterative clustering of neurons and neural Progenitors

To further analyse neurons and neural progenitors, we subset the cells annotated as NP and neurons and generated new Seurat objects and integrated objects as described above. We annotated the cells based on the following markers; *sst1*.*1* (KA’), *urp1* (KA”), slc6a5/*nkx1*.*2lb* (V2s), *olig2*/*sox10* (oligodendrocyte precursor cells, OPC), *zic1*/*zic2a* (*Zic*^*+*^), *nkx2*.*9*/*sulf1* (lateral floor plat, LFP), *fev*/*lmx1bb*/*tph2* (intraspinal serotonergic neurons, ISN), *fabp7a*/*gfap*/*foxj1a* (ependymoradial glia, ERG), *neurod4*/*neurog1* (neural precursor cell, NPC), *slc17a6a*/*slc17a6b* (glutamatergic inhibitory neurons, GLU IN), *foxn4*/*vsx1* (V2 precursor, V2-pre), *mki67*/*sox2*/*sox19a*/*her4* (Neural Progenitors, NP), *isl1*/*isl2a* (motor neuron, MN), *twist1b* (mesodermal cells, MC) and *epcam* (keratinocyte cells, KC). The top markers genes and top transcription factors (obtained from AnimalTFDB (v3.0)) (Hu et al., 2019) were identified from markers genes identified by FindAllMarkers function.

Similarly, we subset ISN cells identified above and re-analysed and integrated them as described above. Then, we identified ISN precursor cells (*fev, lmx1bb*) and ISN cells (*fev* and *tph2*)

### Temporal analyses (pseudotime) of LFP, ISN and V3 cells

To generate the cell trajectory for LFP and ISN cells or V3 cells, we used monocle (version 2) (Cao et al., 2019). The counts and metadata from Seurat objects were used to generate input for monocle, estimateSizefactors, estimateDispersions, detectGenes, differentialGeneTest functions were used sequentially for normalization, estimating dispersion, and selecting genes for ordering. Then, dimensional reduction (reduceDimension) and cell orderings (orderCells) were done. We used a heatmap to visualize the changed genes (qval < 0.00001) based on pseudotime and the num_clusters=2

To perform cell trajectory between LFP and V3 cells, we used LFP and V3 cells and removed LFP_2 (late stage) and MN clusters. We performed the same analyses using monocle. Additionally, we used a publicly available dataset (Scott et al., 2021) (GEO Accession: GSE173350) for LFP and V3 cells. In brief, we performed the same analyses above using Seurat and annotated cells and subset LFP, V3-pre, and V3, using *nkx2*.*2a*/*sim1a* to select V3. Then, we used monocle to generate cell trajectory and pseudotime as described above.

### GO and KEGG pathway analyses

We used clusterProfiler (Wu et al., 2021) package and gene list identified by Seurat or monocle to find enriched GO and KEGGs. Terms with a qvalue < 0.05 were considered statistically significant.

### *In situ* hybridization, immunohistochemistry, sectioning, and cell counting Hybridization chain reaction RNA fluorescent ISH (HCR RNA-FISH)

To generate the *urp1* RNA probe, the sequence of *urp1* was amplified by PCR (The sequences of the oligonucleotides can be found in Table 1 of the supplementary materials).

A 476 bp PCR product was cloned into the pCR®II-TOPO® vector. The plasmid was linearized using the restriction enzyme NotI and the antisense RNA probe was generated using the SP6 RNA polymerase and digoxigenin for labelling (Roche). Whole mount *in situ* hybridization was performed as described previously (Yang et al., 2010).

Hybridization chain reaction RNA fluorescent ISH (HCR RNA-FISH) was performed according to the protocol for whole-mount zebrafish embryos and larvae (Molecular Instruments, Inc.). The *tph2* RNA probe for detecting ISNs was used at a concentration of 16 nM, while the HCR amplifier h1 and h2 (B5-546) were used at a concentration of 6 nM.

The GFP-Booster ATTO488 (1:500, ChromoTek & Proteintech) was used to amplify the fluorescence signal in GFP transgenic larvae.

The head of stained embryos and larvae was cut off manually, and the embryos were embedded laterally in Aqua-Poly/Mount (Polysciences).

### Morpholino knockdown

Injections were performed using a gas-driven microinjector (Tritech Research) as previously described (Muller et al., 1999). Antisense morpholino directed against *nkx2*.*9* was designed by GeneTools. A standard control morpholino was used as a control. The morpholino was resuspended in water with 0.1% phenol red at 0.5 mM. A volume of approximately 1 nl was measured on a calibration micrometre and injected into the yolk of one to two cell-stage embryos. Uninjected embryos and larvae and embryos and larvae injected with standard morpholino (0.5 mM) served as controls. Sequences of the morpholino can be found in Table 1 of the supplementary materials.

### Cell counting

To examine the effect of morpholino injection on the number of cells expressing *urp1*, the stained cells were counted from the 8^th^ to the 16^th^ somite. In the case of *tph2*-expressing cells, all fluorescent intraspinal cells that could be found over the span of five somites above the yolk extension at 3 and 4 dpf were considered. The same procedure was used to count *tph2*^*+*^ cells at 3 dpf in the Notch inhibition experiment.

### LY411575 treatment

LY411575 (Fauq et al., 2007; Sigma Aldrich, St. Louis, MO, USA) was reconstituted with dimethylsulfoxide (DMSO) to a stock concentration of 25 mM. Embryos were incubated in embryonic medium containing 10 µM LY411575, 0.04% DMSO or 0.04% DMSO alone from 48-72 hpf and then washed with embryonic medium containing 0.04% DMSO (Gerber et al., 2019). After treatment, the embryos were fixed with 4%PFA for 2 hours and transferred to 100% methanol at -20 C°.

### Real-Time Quantitative PCR

Zebrafish embryos were collected and manually beheaded. Total RNA from the spinal cord was isolated using Trizol (Life Technology, ThermoFisher Scientific, Darmstadt, Germany) Seventy embryos were used for each condition. Based on the manufacturer’s instructions, the first-strand cDNA was synthesized from 2 µg of total RNA with the Maxima First-Strand cDNA synthesis kit (ThermoFisher Scientific, Darmstadt, Germany). A StepOnePlus Real-Time PCR system (Applied Biosystems, ThermoFisher Scientific, Darmstadt, Germany) and SYBR Green fluorescent dye (Promega, Madison, WI, USA) were used. Expression levels were normalized using β-actin (Zhang et al., 2021). The relative levels of mRNAs were calculated using the 2−ΔΔCT method. Experiments were performed with at least 3 technical replicates. RT-qPCR primer sequences found in Table 1 of the supplementary materials.

### Statistical Analysis

To compare the number of cells expressing the gene of interest between uninjected control or standard control morpholino-injected and morpholino-injected fish, a Welch’s *t*-test was performed using GraphPad Prism 9. Statistical analysis of RT-qPCR results was performed using the unpaired two-tailed Student’s *t*-test. Significance was set at the level of 5 %, meaning that a *p*-value higher or equal to 0.05 represents no significance (n.s.), whereas *p*-values smaller than 0.05 show significant results. The significance level is graded with the number of asterisks: * for *p*<0.05, ** for *p*<0.01, *** for *p*<0.001 and **** for *p*<0.0001.

### Imaging

Images of CISH on zebrafish embryos were acquired with a Leica DM5000B compound microscope with a HC PL APO 63×/1.40-0.60 OIL objective.

For imaging zebrafish larvae stained by HCR RNA-FISH, z-stacks of 0.5 µm intervals were obtained through a TCS SP5 confocal microscope with a HCX APO L 63×/1.2 W U-V-I objective. To obtain single-cell resolution images in control and LY411175 treated group, SP8 inverted confocal microscope (Leica) with a HC PL APO 93×/1.30 GLYC CORR STED WHITE was used.

Fiji/ImageJ was utilized to process images, e.g., for generating Z-projects and adjusting brightness or contrast.

## Figures and figure legends

**Figure-S1: Integration of the datasets and co-expression of *gfp* and *sox1a*** (A) UMAP shows the cell clusters identified at resolution 1. (B) the distribution of cells based on the developmental stages.

**Figure-S3: Enrichment of various cell types at distinct stages of development**. (A) UMAPs show integrated datasets split by developmental stages (from Figure-2A).

**Figure-S4: LFP is the precursor of ISN cells**. (A) Dot plot showing the top marker genes for each cell type. (B) Distribution of cells from each developmental stage on the cell pathway. (C) Pseudotime is shown on cell trajectory. (D) Key transcription factors for LFP and ISN show the cell pathway.

**Figure-S5: LFP is the precursor of both V3 and ISNs**. (A) UMAP shows cell types annotated from (Scott et al., 2021). (B) Feature Plot showing the expression of ISN-pre and ISN markers. (C) The analyses of V3 interneurons from Scott et al. 2021 and their subtypes (left), and the expression of selected marker genes shown on Feature Plots (right). (D) Feature Plots showing marker genes (related to Figure-5A). (E) The integration of LFP, ISN, and V3 cells of the *olig2:eGFP* (Scott et al., 2021) and *sox1a:eGFP* datasets (current study) and the samples are colour-coded. (F) The cell trajectory of LFP, ISN and V3 cells. (G) Pseudotime shown on the cell trajectory. **Note 1**: 24 hpf, 36 hpf, 48 hpf sequencing data are from *olig2:eGFP*^+^ cells (Scott et al., 2021) 1 dpf, 2 dpf, 3 dpf, 5 dpf data are from *sox1a:eGFP*^+^ cells (present work). **Note 2:** CSF-cN1 and CFS-cN2 correspond to KA’ and KA” cells respectively, which we used in this work (Yang et al., 2010).

## Supplementary tables

**Table 1: All_Cells_CellType_Cluster_markers.xlsx**

The top marker genes were identified by using FindAllMarkers from Seurat, using the main cell types as Idents. Related to Figure-1A, B and C.

**Table 2: Sox1a_data_Cell_label_Cluster_markers**

The top marker genes were identified by using FindAllMarkers from Seurat, using the annotated cell clusters as Idents. Related to Figure-2A, B, and C.

**Table 3: ISN_sub-cluster.mrkrs.xlsx**

The top marker genes were identified by using FindAllMarkers from Seurat, using the sub-clusters from the ISN population as Idents. Related to Figure-3C, and D.

**Table 4: LFP_ISN_ISN_pre_all.mrkrs.xlsx**

The top marker genes were identified by using FindAllMarkers from Seurat, using the clusters from the LFP, ISN_pre, and ISN population as Idents. Related to Figure-3C, and D.

**Table 5: LFP ISN_monocle2_trajectory_Sig_gene_Heatmap_Gene_clusters2.xlsx**

The marker genes were identified by differentialGeneTest from monocle and used to generate a pseudotime-dependent heatmap. Related to Figure-4C.

**Table 6: Go_BP_By_Time_sig_genes_heatmap.xlsx**

The GO Biological Process enriched by ClusterProfiler using genes from Table 5. Related to Figure-5D.

**Table 7: KEGG_By_Time_sig_genes_heatmap.xlsx**

The KEGG Pathways enriched by ClusterProfiler using genes from Table 5. Related to Figure-5E.

**Table 8: Olig2_V3_CellType_markers.xlsx**

The top marker genes were identified by using FindAllMarkers from Seurat, using the annotated cell clusters as Idents. Related to Figure-5A.

**Table 9: Sox1a_Olig2_LFP_V3_ISN_Integrated_CellType_markers.xlsx**

The top marker genes were identified by using FindAllMarkers from Seurat, using the annotated cell clusters as Idents. Related to Figure-5C.

**Table 10: LFP_ISN_V3_trajectory_BEAM_Heatmap_Gene_Cluster.xlsx**

The marker genes were identified by BEAM from monocle and used to generate a pseudotime-dependent heatmap. Related to Figure-5E.

## Supplementary materials

Table 1: Oligos, primers and morpholino sequences

---

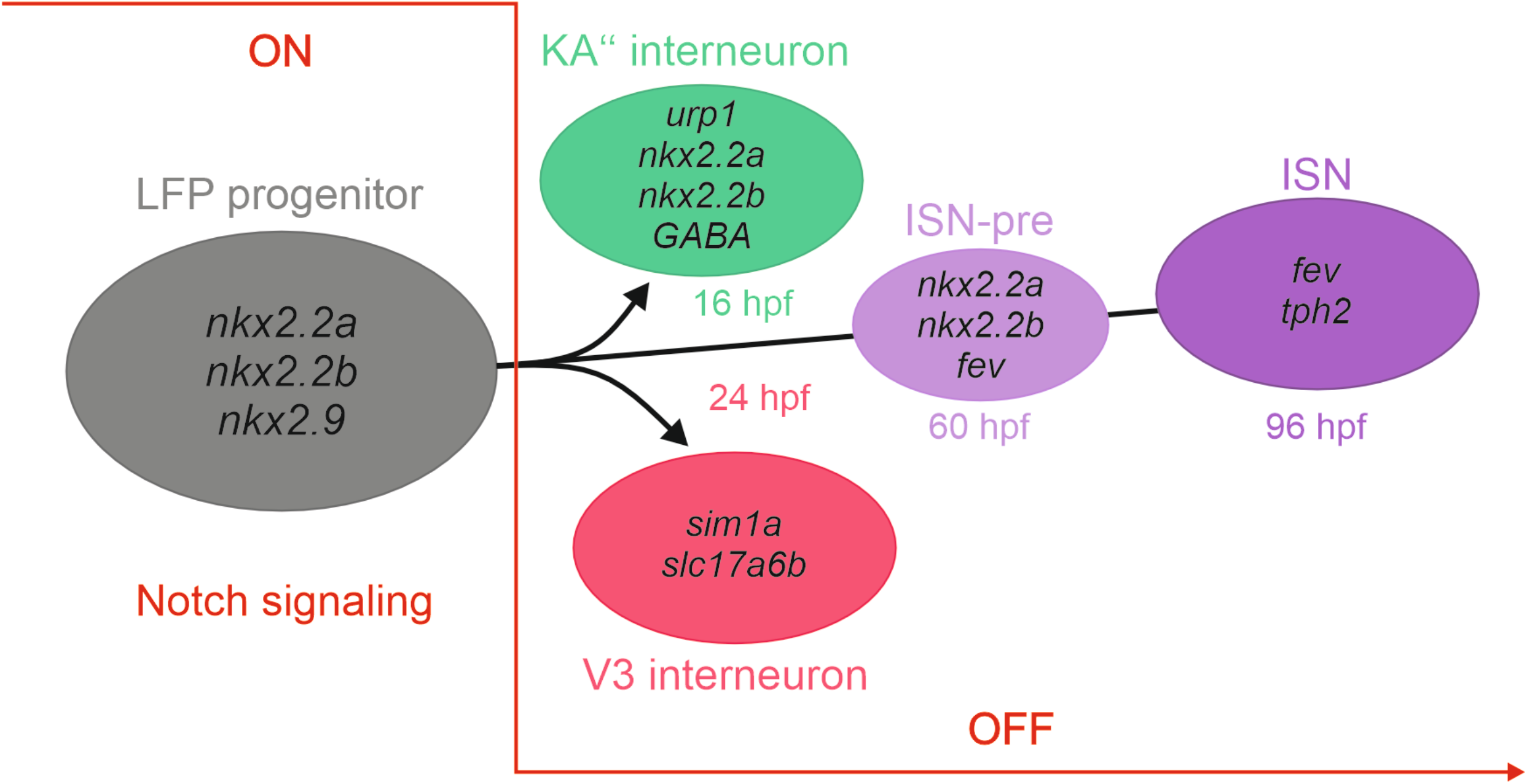

## References

Alaynick, W. A., Jessell, T. M. and Pfaff, S. L. (2011). SnapShot: spinal cord development. Cell 146, 178–178 e171.

Alestrom, P., D’Angelo, L., Midtlyng, P. J., Schorderet, D. F., Schulte-Merker, S., Sohm, F. and Warner, S. (2020). Zebrafish: Housing and husbandry recommendations. Lab Anim 54, 213–224.

Andrews, M. G., Kong, J., Novitch, B. G. and Butler, S. J. (2019). New perspectives on the mechanisms establishing the dorsal-ventral axis of the spinal cord. Curr Top Dev Biol 132, 417–450.

Andrzejczuk, L. A., Banerjee, S., England, S. J., Voufo, C., Kamara, K. and Lewis, K. E. (2018). Tal1, Gata2a, and Gata3 Have Distinct Functions in the Development of V2b and Cerebrospinal Fluid-Contacting KA Spinal Neurons. Front Neurosci 12, 170.

Armant, O., Marz, M., Schmidt, R., Ferg, M., Diotel, N., Ertzer, R., Bryne, J. C., Yang, L., Baader, I., Reischl, M., et al. (2013). Genome-wide, whole mount in situ analysis of transcriptional regulators in zebrafish embryos. Dev Biol 380, 351–362.

Barreiro-Iglesias, A., Mysiak, K. S., Scott, A. L., Reimer, M. M., Yang, Y., Becker, C. G. and Becker, T. (2015). Serotonin Promotes Development and Regeneration of Spinal Motor Neurons in Zebrafish. Cell Rep 13, 924–932.

Batista, M. F., Jacobstein, J. and Lewis, K. E. (2008). Zebrafish V2 cells develop into excitatory CiD and Notch signalling dependent inhibitory VeLD interneurons. Dev Biol 322, 263–275.

Bello-Rojas, S. and Bagnall, M. W. (2022). Clonally related, Notch-differentiated spinal neurons integrate into distinct circuits. Elife 11.

Bill, B. R., Petzold, A. M., Clark, K. J., Schimmenti, L. A. and Ekker, S. C. (2009). A primer for morpholino use in zebrafish. Zebrafish 6, 69–77.

Cao, J., Spielmann, M., Qiu, X., Huang, X., Ibrahim, D. M., Hill, A. J., Zhang, F., Mundlos, S., Christiansen, L., Steemers, F. J., et al. (2019). The single-cell transcriptional landscape of mammalian organogenesis. Nature 566, 496–502.

Cosacak, M. I., Bhattarai, P. and Kizil, C. (2020). Protocol for Dissection and Dissociation of Zebrafish Telencephalon for Single-Cell Sequencing. STAR Protoc 1, 100042.

Cosacak, M. I., Bhattarai, P., Reinhardt, S., Petzold, A., Dahl, A., Zhang, Y. and Kizil, C. (2019). Single-Cell Transcriptomics Analyses of Neural Stem Cell Heterogeneity and Contextual Plasticity in a Zebrafish Brain Model of Amyloid Toxicity. Cell Rep 27, 1307–1318 e1303.

Craven, S. E., Lim, K. C., Ye, W., Engel, J. D., de Sauvage, F. and Rosenthal, A. (2004). Gata2 specifies serotonergic neurons downstream of sonic hedgehog. Development 131, 1165–1173.

Delile, J., Rayon, T., Melchionda, M., Edwards, A., Briscoe, J. and Sagner, A. (2019). Single cell transcriptomics reveals spatial and temporal dynamics of gene expression in the developing mouse spinal cord. Development 146.

Deneris, E. and Gaspar, P. (2018). Serotonin neuron development: shaping molecular and structural identities. Wiley Interdiscip Rev Dev Biol 7.

Deneris, E. S. and Wyler, S. C. (2012). Serotonergic transcriptional networks and potential importance to mental health. Nat Neurosci 15, 519–527.

Dessaud, E., McMahon, A. P. and Briscoe, J. (2008). Pattern formation in the vertebrate neural tube: a sonic hedgehog morphogen-regulated transcriptional network. Development 135, 2489–2503.

Fauq, A. H., Simpson, K., Maharvi, G. M., Golde, T. and Das, P. (2007). A multigram chemical synthesis of the gamma-secretase inhibitor LY411575 and its diastereoisomers. Bioorg Med Chem Lett 17, 6392–6395.

Ferg, M., Armant, O., Yang, L., Dickmeis, T., Rastegar, S. and Strahle, U. (2014). Gene transcription in the zebrafish embryo: regulators and networks. Brief Funct Genomics 13, 131–143.

Gabriel, J. P., Mahmood, R., Kyriakatos, A., Soll, I., Hauptmann, G., Calabrese, R. L. and El Manira, A. (2009). Serotonergic modulation of locomotion in zebrafish: endogenous release and synaptic mechanisms. J Neurosci 29, 10387–10395.

Gerber, V., Yang, L., Takamiya, M., Ribes, V., Gourain, V., Peravali, R., Stegmaier, J., Mikut, R., Reischl, M., Ferg, M., et al. (2019). The HMG box transcription factors Sox1a and Sox1b specify a new class of glycinergic interneuron in the spinal cord of zebrafish embryos. Development 146.

Goulding, M. (2009). Circuits controlling vertebrate locomotion: moving in a new direction. Nat Rev Neurosci 10, 507–518.

Hao, Y., Hao, S., Andersen-Nissen, E., Mauck, W. M., 3rd, Zheng, S., Butler, A., Lee, M. J., Wilk, A. J., Darby, C., Zager, M., et al. (2021). Integrated analysis of multimodal single-cell data. Cell 184, 3573–3587 e3529.

Hendricks, T., Francis, N., Fyodorov, D. and Deneris, E. S. (1999). The ETS domain factor Pet-1 is an early and precise marker of central serotonin neurons and interacts with a conserved element in serotonergic genes. J Neurosci 19, 10348–10356.

Hendricks, T. J., Fyodorov, D. V., Wegman, L. J., Lelutiu, N. B., Pehek, E. A., Yamamoto, B., Silver, J., Weeber, E. J., Sweatt, J. D. and Deneris, E. S. (2003). Pet-1 ETS gene plays a critical role in 5-HT neuron development and is required for normal anxiety-like and aggressive behavior. Neuron 37, 233–247.

Hu, H., Miao, Y. R., Jia, L. H., Yu, Q. Y., Zhang, Q. and Guo, A. Y. (2019). AnimalTFDB 3.0: a comprehensive resource for annotation and prediction of animal transcription factors. Nucleic Acids Res 47, D33–D38.

Huang, C. X., Zhao, Y., Mao, J., Wang, Z., Xu, L., Cheng, J., Guan, N. N. and Song, J. (2021). An injury-induced serotonergic neuron subpopulation contributes to axon regrowth and function restoration after spinal cord injury in zebrafish. Nat Commun 12, 7093.

Huang, P., Xiong, F., Megason, S. G. and Schier, A. F. (2012). Attenuation of Notch and Hedgehog signaling is required for fate specification in the spinal cord. PLoS Genet 8, e1002762.

Jacobs, C. T. and Huang, P. (2019). Notch signalling maintains Hedgehog responsiveness via a Gli-dependent mechanism during spinal cord patterning in zebrafish. Elife 8.

Jacobs, C. T., Kejriwal, A., Kocha, K. M., Jin, K. Y. and Huang, P. (2022). Temporal cell fate determination in the spinal cord is mediated by the duration of Notch signalling. Dev Biol 489, 1–13.

Karunaratne, A., Hargrave, M., Poh, A. and Yamada, T. (2002). GATA proteins identify a novel ventral interneuron subclass in the developing chick spinal cord. Dev Biol 249, 30–43.

Kimmel, C. B., Ballard, W. W., Kimmel, S. R., Ullmann, B. and Schilling, T. F. (1995). Stages of embryonic development of the zebrafish. Dev Dyn 203, 253–310.

Kimura, Y., Satou, C. and Higashijima, S. (2008). V2a and V2b neurons are generated by the final divisions of pair-producing progenitors in the zebrafish spinal cord. Development 135, 3001–3005.

Krueger, K. C. and Deneris, E. S. (2008). Serotonergic transcription of human FEV reveals direct GATA factor interactions and fate of Pet-1-deficient serotonin neuron precursors. J Neurosci 28, 12748–12758.

Lai, H. C., Seal, R. P. and Johnson, J. E. (2016). Making sense out of spinal cord somatosensory development. Development 143, 3434–3448.

Le Dreau, G. and Marti, E. (2012). Dorsal-ventral patterning of the neural tube: a tale of three signals. Dev Neurobiol 72, 1471–1481.

Li, S., Misra, K., Matise, M. P. and Xiang, M. (2005). Foxn4 acts synergistically with Mash1 to specify subtype identity of V2 interneurons in the spinal cord. Proc Natl Acad Sci U S A 102, 10688–10693.

Lillesaar, C., Tannhauser, B., Stigloher, C., Kremmer, E. and Bally-Cuif, L. (2007). The serotonergic phenotype is acquired by converging genetic mechanisms within the zebrafish central nervous system. Dev Dyn 236, 1072–1084.

Lu, D. C., Niu, T. and Alaynick, W. A. (2015). Molecular and cellular development of spinal cord locomotor circuitry. Front Mol Neurosci 8, 25.

McMahon, A. P. (2000). Neural patterning: the role of Nkx genes in the ventral spinal cord. Genes Dev 14, 2261–2264.

Montgomery, J. E., Wahlstrom-Helgren, S., Wiggin, T. D., Corwin, B. M., Lillesaar, C. and Masino, M. A. (2018). Intraspinal serotonergic signaling suppresses locomotor activity in larval zebrafish. Dev Neurobiol.

Montgomery, J. E., Wiggin, T. D., Rivera-Perez, L. M., Lillesaar, C. and Masino, M. A. (2016). Intraspinal serotonergic neurons consist of two, temporally distinct populations in developing zebrafish. Dev Neurobiol 76, 673–687.

Muller, F., Chang, B., Albert, S., Fischer, N., Tora, L. and Strahle, U. (1999). Intronic enhancers control expression of zebrafish sonic hedgehog in floor plate and notochord. Development 126, 2103–2116.

Odenthal, J., van Eeden, F. J., Haffter, P., Ingham, P. W. and Nusslein-Volhard, C. (2000). Two distinct cell populations in the floor plate of the zebrafish are induced by different pathways. Dev Biol 219, 350–363.

Okuda, Y., Ogura, E., Kondoh, H. and Kamachi, Y. (2010). B1 SOX coordinate cell specification with patterning and morphogenesis in the early zebrafish embryo. PLoS Genet 6, e1000936.

Panayi, H., Panayiotou, E., Orford, M., Genethliou, N., Mean, R., Lapathitis, G., Li, S., Xiang, M., Kessaris, N., Richardson, W. D., et al. (2010). Sox1 is required for the specification of a novel p2-derived interneuron subtype in the mouse ventral spinal cord. J Neurosci 30, 12274–12280.

Pattyn, A., Vallstedt, A., Dias, J. M., Samad, O. A., Krumlauf, R., Rijli, F. M., Brunet, J. F. and Ericson, J. (2003). Coordinated temporal and spatial control of motor neuron and serotonergic neuron generation from a common pool of CNS progenitors. Genes Dev 17, 729–737.

Reiter, F., Wienerroither, S. and Stark, A. (2017). Combinatorial function of transcription factors and cofactors. Curr Opin Genet Dev 43, 73–81.

Sagner, A. and Briscoe, J. (2019). Establishing neuronal diversity in the spinal cord: a time and a place. Development 146.

Scott, K., O’Rourke, R., Winkler, C. C., Kearns, C. A. and Appel, B. (2021). Temporal single-cell transcriptomes of zebrafish spinal cord pMN progenitors reveal distinct neuronal and glial progenitor populations. Dev Biol 479, 37–50.

Scott, M. M., Krueger, K. C. and Deneris, E. S. (2005). A differentially autoregulated Pet-1 enhancer region is a critical target of the transcriptional cascade that governs serotonin neuron development. J Neurosci 25, 2628–2636.

Smith, E., Hargrave, M., Yamada, T., Begley, C. G. and Little, M. H. (2002). Coexpression of SCL and GATA3 in the V2 interneurons of the developing mouse spinal cord. Dev Dyn 224, 231–237.

Strahle, U., Scholz, S., Geisler, R., Greiner, P., Hollert, H., Rastegar, S., Schumacher, A., Selderslaghs, I., Weiss, C., Witters, H., et al. (2012). Zebrafish embryos as an alternative to animal experiments--a commentary on the definition of the onset of protected life stages in animal welfare regulations. Reprod Toxicol 33, 128–132.

Westerfield, M. (2000). The Zebrafish Book -A Guide for the Laboratory use of Zebrafish (Brachydanio rerio). University of Oregon Press.

Wu, T., Hu, E., Xu, S., Chen, M., Guo, P., Dai, Z., Feng, T., Zhou, L., Tang, W., Zhan, L., et al. (2021). clusterProfiler 4.0: A universal enrichment tool for interpreting omics data. Innovation (Camb) 2, 100141.

Wyler, S. C., Spencer, W. C., Green, N. H., Rood, B. D., Crawford, L., Craige, C., Gresch, P., McMahon, D. G., Beck, S. G. and Deneris, E. (2016). Pet-1 Switches Transcriptional Targets Postnatally to Regulate Maturation of Serotonin Neuron Excitability. J Neurosci 36, 1758–1774.

Xu, J., Srinivas, B. P., Tay, S. Y., Mak, A., Yu, X., Lee, S. G., Yang, H., Govindarajan, K. R., Leong, B., Bourque, G., et al. (2006). Genomewide expression profiling in the zebrafish embryo identifies target genes regulated by Hedgehog signaling during vertebrate development. Genetics 174, 735–752.

Yang, L., Rastegar, S. and Strahle, U. (2010). Regulatory interactions specifying Kolmer-Agduhr interneurons. Development 137, 2713–2722.

Yang, L., Wang, F. and Strahle, U. (2020). The Genetic Programs Specifying Kolmer-Agduhr Interneurons. Front Neurosci 14, 577879.

Zhang, G., Lubke, L., Chen, F., Beil, T., Takamiya, M., Diotel, N., Strahle, U. and Rastegar, S. (2021). Neuron-Radial Glial Cell Communication via BMP/Id1 Signaling Is Key to Long-Term Maintenance of the Regenerative Capacity of the Adult Zebrafish Telencephalon. Cells 10.

Zheng, G. X., Terry, J. M., Belgrader, P., Ryvkin, P., Bent, Z. W., Wilson, R., Ziraldo, S. B., Wheeler, T. D., McDermott, G. P., Zhu, J., et al. (2017). Massively parallel digital transcriptional profiling of single cells. Nat Commun 8, 14049.

Zhou, Y., Yamamoto, M. and Engel, J. D. (2000). GATA2 is required for the generation of V2 interneurons. Development 127, 3829–3838.

